# Harvesting information from ultra-short ancient DNA sequences

**DOI:** 10.1101/319277

**Authors:** Cesare de Filippo, Matthias Meyer, Kay Prüfer

## Abstract

The study of ancient DNA is hampered by degradation, resulting in short DNA fragments. Advances in laboratory methods have made it possible to retrieve short DNA fragments, thereby improving access to DNA preserved in highly degraded, ancient material. However, such material contains large amounts of microbial contamination in addition to DNA fragments from the ancient organism. The resulting mixture of sequences constitute a challenge for computational analysis, since microbial sequences are hard to distinguish from the ancient sequences of interest, especially when they are short. Here, we develop a method to quantify spurious alignments based on the presence or absence of rare variants. We find that spurious alignments are enriched for mismatches and insertion/deletion differences and lack substitution patterns typical of ancient DNA. The impact of spurious alignments can be reduced by filtering on these features and by imposing a sample-specific minimum length cutoff. We apply this approach to sequences from the ~430,000 year-old Sima de los Huesos hominin remains, which contain particularly short DNA fragments, and increase the amount of usable sequence data by 17-150%. This allows us to place a third specimen from the site on the Neandertal lineage. Our method maximizes the sequence data amenable to genetic analysis from highly degraded ancient material and avoids pitfalls that are associated with the analysis of ultra-short DNA sequences.

## Background

After its death, the DNA of an organism inevitably degrades into short DNA fragments [1, 2]. Laboratory methods have been developed that specifically aim at retrieving these fragments from ancient biological material [3–5] and transforming them efficiently into library molecules for high-throughput sequencing [6]. These developments have enabled researchers to study DNA sequences from increasingly older samples. One notable example are the remains from Sima de los Huesos in Spain that constitute, with an age of over 400,000 years, the by far oldest hominin material to date that yielded ancient DNA sequences [7, 8]. Owing to their great age, the vast majority of hominin DNA fragments that can be extracted from the Sima de los Huesos remains are shorter than 45 bp [7].

In addition to the extreme state of DNA fragmentation, the analysis of sequences from highly degraded material is hampered by the large number of extraneous DNA fragments originating from microorganisms that decomposed the remains of the source organism after its death [9–12]. In the case of Sima de los Huesos [8] and many other ancient skeletal remains, microbial DNA constitutes more than 99% of the DNA that can be recovered and sequenced. Contaminant sequences are typically differentiated from those that stem from the source organism by aligning all sequences to a related reference genome and retaining only those that produce alignments with not more than a pre-defined number of differences [13, 14]. However, unrelated sequences can align by chance and the probability of such spurious alignments increases with decreasing sequence length [15]. This issue is expected to affect particularly the analysis of sequences from highly fragmented material.

To minimize the effect of spuriously aligning sequences on downstream analyses, previous studies employed sequence length cutoffs that have been gauged by a variety of methods. Green et al. [16] used specific alignment software to analyze the distribution of alignment scores at various sequence lengths. This distribution was found to be distinctly bimodal at longer lengths, as expected from a mixture of related and unrelated sequence alignments, while bimodality was not observed at shorter lengths. Setting a length cutoff that preserves the bimodal distribution can thus be used to limit the fraction of spurious alignments. Cutoffs have also been determined by testing at which lengths mammoth sequences yielded equally good alignments to other mammalian taxa [10, 13], horse sequences aligned equally well to the chicken genome [17], mammoth and ancient bovine sequences aligned to a database of concatenated bacterial genomes [18], or fragmented bacterial genomes aligned to the human reference [19]. While these methods have been sufficient to determine approximate cutoffs, they do not provide an estimate of the fraction of spurious alignments. We also note that microbial genomes in public databases may present a poor proxy for the microbial sequence diversity found in real sequence data from ancient remains. The validity of these approaches is therefore hard to judge.

More recently, Meyer et al. [8] used a different approach to determine sequence length cutoffs for the analysis of nuclear DNA sequences from the Sima de los Huesos samples. Using sequence variants that are unique to the human reference genome, as determined by comparison to known variation from human resequencing studies and the genomes of non-human primates, they counted the fraction of sequences that match the reference-specific variant. These variants are rare and are expected to be largely absent in other hominin genomes. In contrast, spuriously aligned sequences will match the reference genome by chance, independent of how frequent the reference genomes’ variants are in the human population. Since no matches to the reference-specific variant was observed for sequences of at least 35 bp length, this cutoff was deemed sufficient to exclude spurious alignments. However, due to the limited number of unique reference variants (i.e. 11,299) and the small amount of data obtained from the Sima de los Huesos remains (less than 0.001-fold genomic coverage per sample), only between 4 and 69 sequences formed the basis for this assessment, preventing any fine-scale estimates of the fraction of spurious alignments.

Here we test and extend this approach to allow for the confident estimation of the fraction of spurious alignments across different sequence lengths. We use these estimates to devise sequence length cutoffs that maximize the number of useful sequences and increase the power of phylogenetic analysis. Applying our approach to the Sima de los Huesos samples, we determine that cutoffs shorter than 35 bp are suitable for some of these samples, as long as appropriate filters are put in place. The increase in usable sequences allows us to confidently place one of the Sima de los Huesos samples on the Neandertal lineage that previously yielded inconclusive results.

## Results

### Estimating the fraction of spurious alignments

To allow for fine-scale estimates of the fraction of spurious alignments in small data sets, we changed ~18 million interspersed bases in the human reference genome (see Methods). These artificial mutations were introduced at positions where the human reference, all human genomes sequenced as part of the 1000 Genomes project, two high-coverage archaic human genomes and the chimpanzee genome show the same base. They are thus unlikely to occur in present-day or ancient hominin genomes (probability<0.1%). Spurious alignments, on the other hand, are likely to match the mutated state (Fig. 1A). The alignment parameters used here and in other studies [e.g. 14, 21] limit the fraction of allowed mismatches per alignment to approximately 10% (see Material and Methods), resulting for spuriously aligned sequences in a predicted ~90% match probability for the mutated state and a ~3.3% probability for matching either of the remaining three states (Fig. 1A).

**Fig. 1.**
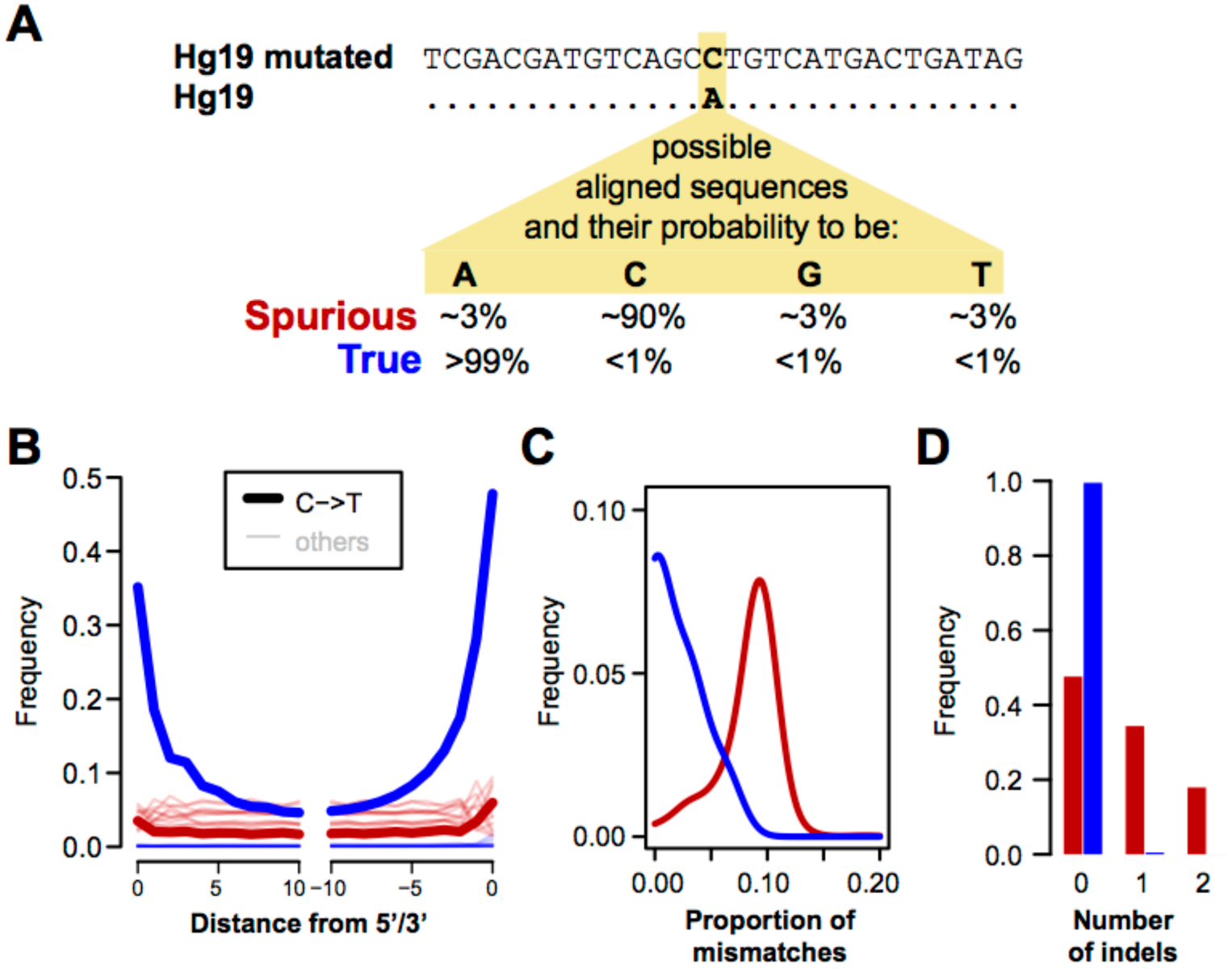
Identification and characterization of spurious and true sequence alignments. **(A)** Schematic illustration of how spurious and true sequence alignments are inferred. The human reference (hg19) is mutated to introduce changes at positions that are not known to vary among present-day humans and other hominins. True hominin sequences (blue) and spuriously aligned microbial sequences (red) are expected to show the reference, the mutated or one of the two other states with the probabilities indicated. **(B)** Frequency of all nucleotide substitutions at each position in Mezmaiskaya 1 sequence alignments. **(C)** Distributions of the proportion of mismatches in Mezmaiskaya 1 alignments. One mismatch was subtracted from all true alignments and those spurious alignments that did not carry the mutated allele. This was done to compensate for the fact that these alignments have to carry a mismatch to the mutated reference genome in order to be identified as such. **(D)** Distributions of the number of indels in Mezmaiskaya 1 alignments.

To test whether these predictions hold, we generated sequences from DNA isolated from the blood sample of a healthy human individual that was fragmented heavily to mimic the size distribution of ancient DNA. We further compiled a dataset consisting of 3,860 bacterial genomes that were cut *in silico* into 7.2 billion unique sequences of 32 bp length (see Materials and Methods). We then mapped both sets of sequences to the mutated reference and counted the fraction of sequences that match the reference state at mutated positions (presumed hominin alignment, henceforth “true alignment”) or any other variant (presumed “spurious alignment”). Of the aligned human sequences, 99.8% were correctly classified as hominin. Out of 15.1 million bacterial sequences that could be aligned to the mutated reference, 96.98% were correctly classified as spurious. If all alignments of bacterial sequences contained the maximal number of allowed mismatches, 3.12% of the sequences would be expected to carry the reference state by chance, which is close to the 3.02% observed. Since these ~3% of misclassified sequences bias the estimated fraction of spurious alignments slightly downward, we corrected our estimates in all subsequent analyses accordingly (see Materials and Methods).

### Characteristics of spurious and true alignments

We next investigated whether spurious and true alignments differ in specific characteristics. For this purpose we aligned sequences from the Mezmaiskaya1 Neandertal [21, 22], a published dataset containing a considerable fraction of ultra-short (<35bp) sequences (Fig. S1) and approximately 9% Neandertal DNA, to the mutated reference. After filtering for mappability (see Methods) and classifying the alignments as described above, we obtained 5.07 million true alignments and 0.92 million spurious alignments.

We first note that true Mezmaiskaya 1 sequence alignments show elevated frequencies of C-to-T substitutions, which occur predominantly at their beginning and ends (Fig. 1B). This pattern is expected for authentic ancient DNA sequences and results from deamination of cytosine to uracil in single-stranded DNA overhangs [23]. In contrast, this pattern is not observed for spurious alignments, where C-to-T substitutions are similar in frequency to other types of substitutions. Second, we find that true alignments carry significantly fewer mismatches on average than spurious alignments (0.018 vs 0.108 per bp; Wilcoxon rank sum test p-value<2.2e-16; see Fig. 1C). The fraction of mismatches in the true alignments is still substantially larger than the genomic divergence between modern humans and Neandertals of <0.002 differences per base pair [22]. However, C-to-T substitutions account for most of this difference (Fig. 1B). Third, true alignments contain fewer insertions/deletions (indels) than spurious alignments (0.5% vs. 52.4% of the alignments, Wilcoxon rank sum test, p-value<2.2e-16) (Fig. 1D). Indels accumulate at a roughly 10-times lower rate than single nucleotide mutations in humans [24] and are therefore expected to be rare in true alignments.

We repeated these analyses using the bacterial and modern human control datasets. Similar to the results from spurious Mezmaiskaya 1 alignments, bacterial alignments are enriched for mismatches (0.092 per bp on average) and indels (76.2% of the alignments), whereas mismatches and indels are rare among modern human control alignments (0.004 per bp and 0.05%, respectively).

### Minimizing the proportion of spurious alignments

We next binned all Mezmaiskaya 1 sequences by length and calculated the fraction of spurious alignments for each bin. As expected, the fraction of spurious alignments increases with decreasing sequence length (Fig. 2). Spurious alignments are rare (<0.3%) in sequences of at least 35 bp length, suggesting that a sequence length cutoff of 35 bp, which was used in several ancient DNA studies (Table S1), is effective in removing the vast majority of spurious alignments for the Mezmaiskaya 1 dataset analyzed here. In fact, even sequences of length 33 bp show a proportion of spurious alignments of less than 1%, indicating that shorter sequences could be included in downstream analyses (Fig. 2).

**Fig. 2.**
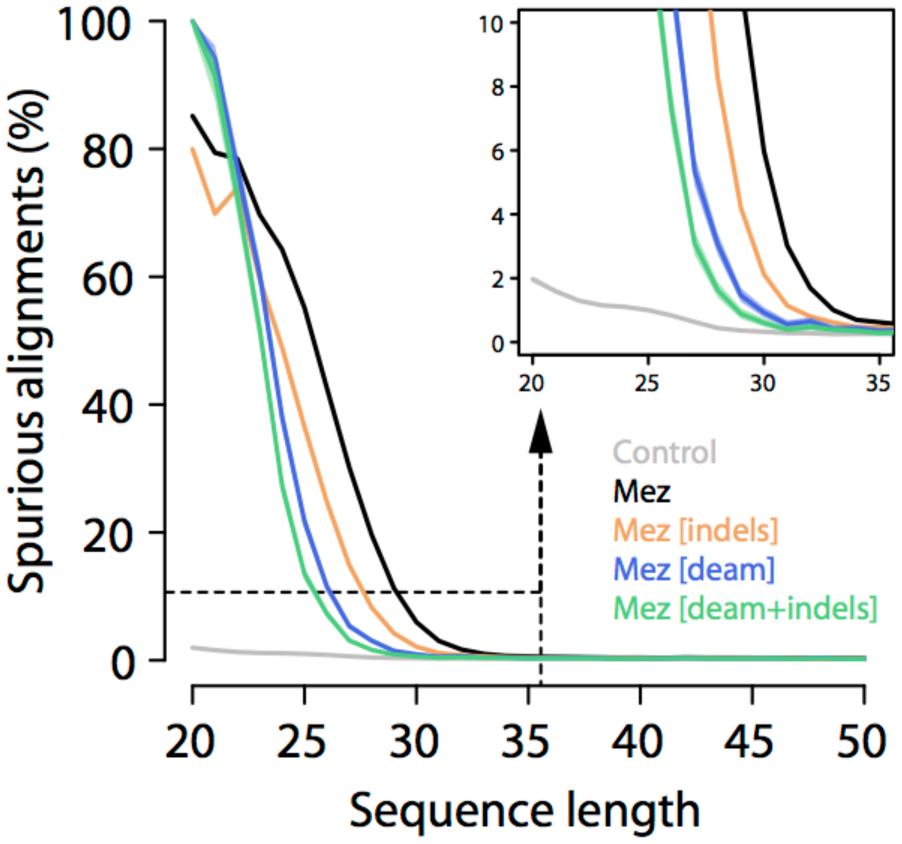
Effect of sequence length, indel and deamination filters on the proportion of spurious alignments. The proportion of spurious alignments in each size bin is shown for all Mezmaiskaya 1 (“Mez”) alignments in black, alignments without indels in orange, alignments with terminal C to T substitutions (“deam”) in blue, and with both filters applied in green. An analysis of modern human sequences (“Control”) without microbial contamination is shown in gray. 95% binomial confidence intervals are as wide as or smaller than the line width.

The previous analysis has shown that spurious alignments lack the elevation of terminal C-to-T substitution frequencies that are typical for ancient DNA and that they contain more indels than true alignments (Fig. 1). Filtering based on these features may thus help to further reduce the fraction of spurious alignments. In agreement with this assumption, we find that restricting the analysis to alignments exhibiting a C to T substitution at either terminus yields less than 1% spurious alignments for length bins as short as 30 bp. It should be noted that this deamination filter is often used to deplete sequence data of human contamination. However, it also removes a large fraction of potentially genuine ancient sequences that were not affected by deamination (~85% of aligned sequences ≥ 35 bp in Mezmaiskaya 1). A less pronounced effect is observed when removing alignments with indels (~1% of aligned sequences ≥ 35 bp in Mezmaiskaya 1), which yields less than 1% spurious alignments in size bins of 32 bp or longer. Combining both filters reduces this number to 29 bp. The reduction of spurious alignments achieved with both filters is also reflected by a decrease in sequence differences to the reference genome (Fig. 2).

We repeated this analysis using our modern human control sample, which should, by design, not produce any spurious alignments. We find that even the shortest length bin yields an estimate for the proportion of spurious alignments of less than 2% (Fig. 2), suggesting that sequencing or mapping errors have little impact on our measure.

### A re-analysis of sequences from Sima de los Huesos

The extremely short DNA sequences that have been retrieved from the Sima de los Huesos remains are an ideal dataset to explore to which extent the choice of sequence filters changes the amount of useful sequence data that can be obtained from very poorly preserved material and the inferences that can be drawn from these data. Appreciable amounts of nuclear DNA sequences are available from four hominin remains from the site [8]. The fraction of hominin DNA varies between 0.02% and 0.18% in these samples when considering sequences of at least 35 bp length. However, the vast majority (>97%) of the human aligned sequences of at least 20 bp length are shorter than this 35 bp cutoff (Fig. S1).

To determine whether at least some of these ultra-short sequences are amenable to analysis, we realigned the data of all four samples to the mutated reference genome and removed alignments that contained indels and those showing no evidence of deamination (see also Fig. S2-S3). The deamination filter is strictly required when working with these data, as a substantial fraction of the hominin sequences is derived from modern human contamination [7, 8]. We then calculated sequence length cutoffs that limit the fraction of spurious alignments to <1% or <10%, henceforth denoted by *L_1%_* and *L_10%_*, respectively.

The four samples yield *L_10%_* cutoffs that range from 27 to 34 bp and decrease with increasing proportions of endogenous DNA (Fig. 3, Table 1). Applying these cutoffs instead of the previously used cutoff of 35 bp would increase the usable data by 17-150%. The more conservative *L_1%_* cutoffs would result in 0-40% more data for three of the four samples. Interestingly, the fourth sample, FemurXIII, yields a *L_1%_* cutoff of 46 bp, suggesting that the often applied cutoff of 35 bp is not always sufficient to limit spurious alignments to low levels. In comparison, sequences from the Mezmaiskaya 1 Neandertal yield an *L_1%_* of 22 bp and do not reach a limit for *L_10%_* (less than 10% of all sequences of at least 20 bp length aligned spuriously). Considering sequences of at least 20 bp for analysis would result in 37% more data compared to a 35 bp length cutoff.

**Fig. 3.**
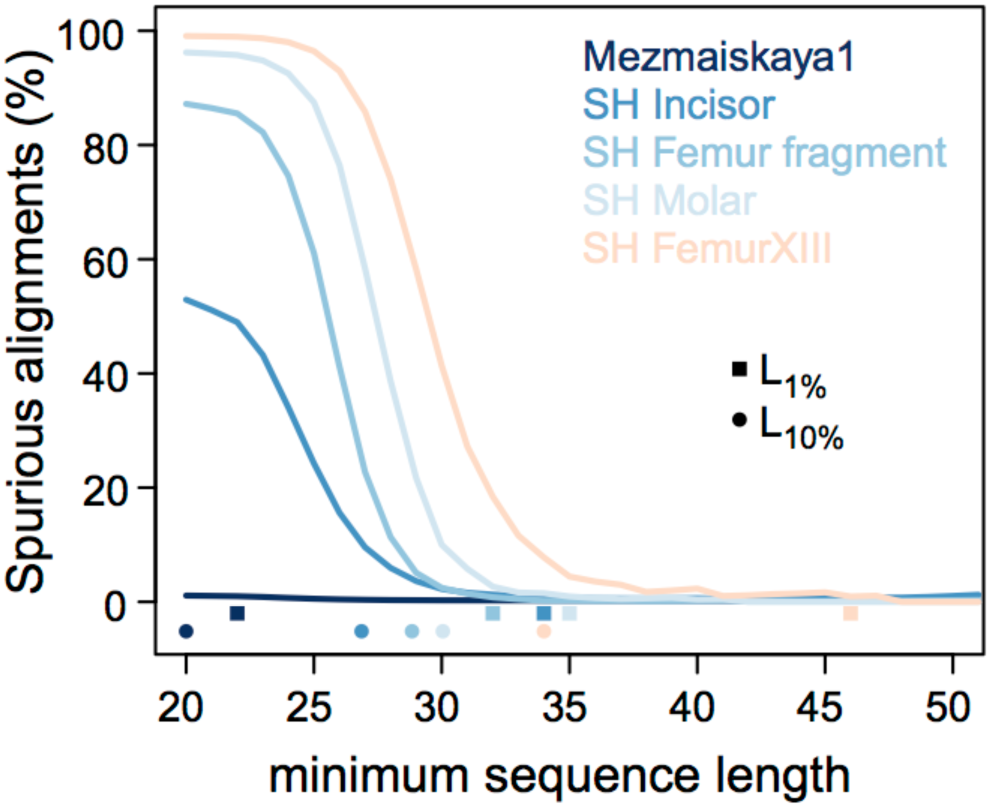
Cumulative proportion of spurious alignments. Squares and dots on the x-axis show the length-cutoffs that guarantee a spurious alignment rate lower than 1% (*L_1%_*) and lower than 10% (*L_10%_*), respectively (see also Table 1). Only sequences with C-to-T changes in the terminal 5’and 3’ positions, and without indels are considered (i.e. the filters “deam+indels” used in Fig. 2).

**Table 1.** Mezmaiskaya and Sima de los Huesos (SH) sequence length cutoffs allowing for less than 1% or 10% spurious alignments.

| Samples | Hominin DNA (%)# | Length cutoff (bp)§ |  | Total number of hominin bases recovered (Mbp) |  |  | Fold-change |  |
| --- | --- | --- | --- | --- | --- | --- | --- | --- |
| | | $L_{1\%}$ | $L_{10\%}$ | 35bp | $L_{1\%}$ | $L_{10\%}$ | $L_{1\%}/35bp$ | $L_{10\%}/35bp$ |
| Mezmaiskaya1 | 8.84 | 22 | 20 | 88.53 | 121.17 | 121.73 | 1.37 | 1.37 |
| SH Femur frag. | 0.11 | 32 | 29 | 0.81 | 1.14 | 1.52 | 1.41 | 1.88 |
| SH Incisor | 0.18 | 34 | 27 | 1.49 | 1.70 | 3.71 | 1.14 | 2.49 |
| SH Molar | 0.03 | 35 | 30 | 0.13 | 0.13 | 0.25 | 1.00 | 1.98 |
| SH FemurXIII | 0.02 | 46 | 34 | 0.15 | 0.03 | 0.18 | 0.23 | 1.17 |

Since present-day human contamination constitutes a challenge for the analysis of archaic human sequences, we also tested whether contamination rates differ when including shorter sequences. We found no significant differences in the estimated contamination compared to the previously used length cutoff of 35 bp (Table S2).

### Improving phylogenetic inferences from limited data

The initial analysis of nuclear DNA sequences from the Sima de los Huesos specimens revealed that two of the specimens (an incisor and a femur fragment) share significantly more derived alleles with the high-coverage genome of a Neandertal than with that of a Denisovan individual [8]. While this result concurred with the fact that the Middle Pleistocene Sima de los Huesos remains were discovered in the western part of the territory inhabited by Neandertals during the Late Pleistocene (Europe and Central Asia), it deviated from the mitochondrial tree [7], which groups the Sima de los Huesos hominins into a clade with Denisovans, who are thought to have inhabited large parts of Asia [25, 26].

To test whether the inclusion of data from shorter nuclear sequences would affect inferences about the phylogenetic position of the Sima de los Huesos specimens, we compared the results of the lineage assignment test (see Material and Methods) obtained by using a 35 bp cutoff, as previously published, and the *L_10%_* cutoffs determined here to all four specimens for which at least 1,000 sequences from putatively deaminated DNA fragments were available (Fig. 4). For the femur fragment and the incisor, the inclusion of additional data strengthens the confidence of the Neandertal lineage assignment, and the significance of the assignment was highest when between 2.5% and 15.6% of spurious alignments were allowed (Fig. S4). This suggests that a spurious alignment proportion of around 10% can be tolerated for this analysis. As previously, one of the other Sima de los Huesos samples (Femur XIII) did not yield sufficient data for a confident lineage assignment and no additional data could be gained by applying the *L_10%_* cutoff. However, the fourth specimen, a molar, shows significantly higher allele sharing with the Neandertal than the Denisovan genome with the *L_10%_* cutoff (Fig. 4, Fisher exact test p-value = 0.005 corrected for multiple testing [27]). Moreover, the percentage of Neandertal-shared derived alleles of the molar (35%) does not significantly differ from the percentages observed for the incisor and the femur fragment (43% and 39%, respectively; all pairwise Fisher’s exact tests p-values > 0.29).

**Fig. 4.**
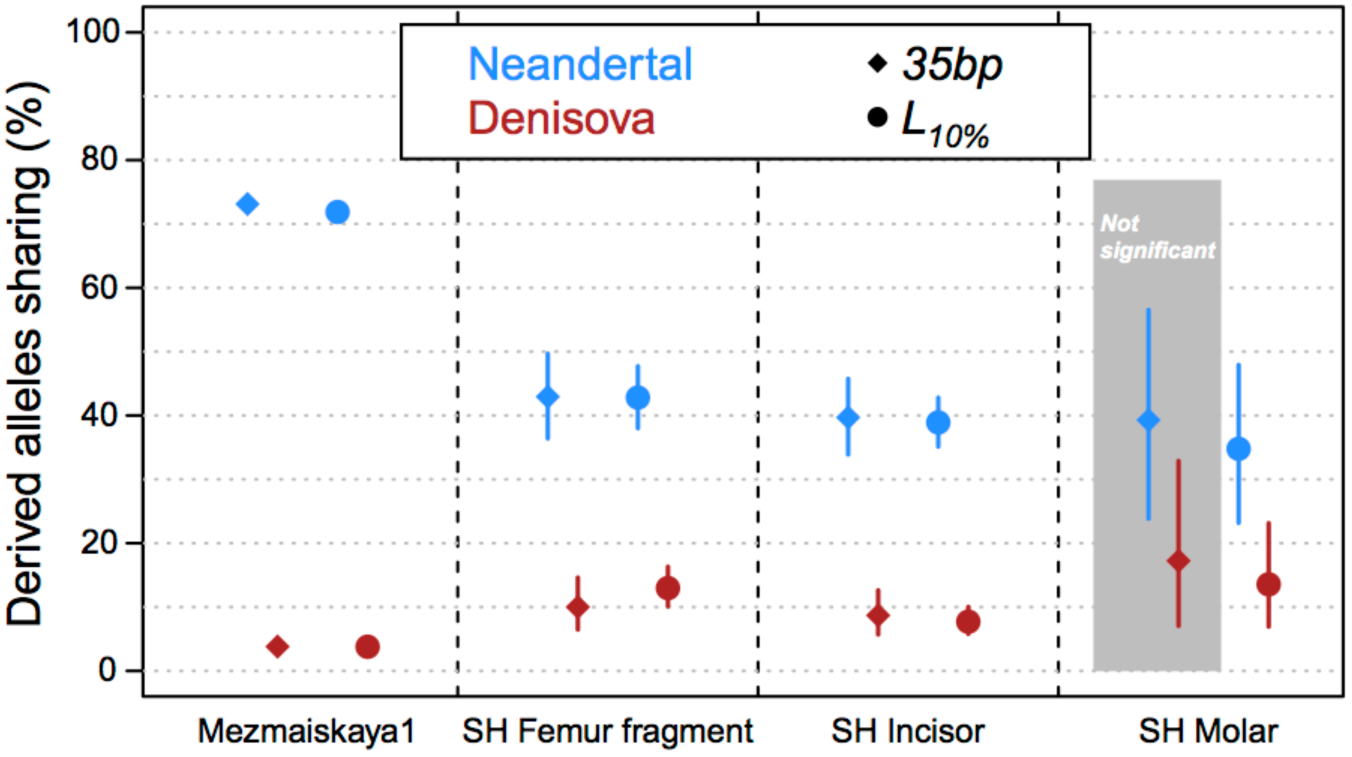
Percentage of derived allele sharing with the Denisovan and Neandertal lineages. The circles and diamonds correspond to the *L_10%_* and *35bp* length cutoffs, respectively. Bars indicate 90% binomial confidence intervals. The difference between Neandertal and Denisovan sharing is statistically significant in all comparisons, except for the SH Molar with the 35bp cutoff (Fisher exact test p-value=0.09).

### Phylogenetic inferences and reference bias

Since the fraction of mismatches in alignments is limited, spuriously aligning sequences are expected to exhibit a strong bias towards showing the human reference allele. This preference for the human reference allele should introduce a bias towards supporting the modern human lineage in the lineage assignment analysis of spurious alignments. In agreement with this expectation, we observe a strong bias towards the human reference allele in misaligning bacterial sequences, which are assigned to the modern human lineage (~33% of the human derived variants shared). A similar signal is also observed for the lineage assignment of Sima de los Huesos when considering size cutoffs that are expected to lead to an overwhelming majority of spurious alignments (Figure S5). While our results with the *L_10%_* cutoffs do not show significant differences to previous, more conservative cutoffs for these samples, we caution that reference bias may affect analyses and needs to be considered before including a higher fraction of spurious alignments.

## Discussion/Conclusions

Experimental procedures have made great strides forward in extracting short ancient DNA fragments [3, 5, 6]. However, the resulting short sequences constitute a challenge for computational processing since unrelated and related sequences cannot easily be distinguished. This has led to the paradoxical situation, in which short DNA fragments that are preserved in highly degraded samples can be made accessible to sequencing, only to be discarded in downstream computational analyses to avoid spurious alignments.

How can shorter sequences be made available for analysis without increasing the fraction of spurious alignments unduly? We have shown here that one answer lies in specific filters that enrich for genuine alignments. By filtering for sequences with evidence for deamination and without insertion/deletion differences to the reference genome, we were able to reduce the fraction of spurious alignments sufficiently to allow for the inclusion of sequences shorter than 35 bp from three Sima de los Huesos samples in phylogenetic analysis. This analysis confirmed that two of the samples originate from early Neandertals and enabled us to place one additional sample, a molar, on the Neandertal lineage. The Neandertal allele sharing of this sample is similar to that of the other two. All three samples could thus originate from a single group of early Neandertal ancestors or relatives thereof.

The highly degraded remains from Sima de los Huesos yielded, arguably, the most challenging dataset in ancient DNA to date, containing a large fraction of ultra-short sequences and a large fraction of sequences from microbial contamination. In light of these difficulties it is encouraging for future work on material with poor DNA preservation that useful genetic information could be recovered from ultra-short sequences of three samples from the site. However, we have to acknowledge that working with such sequences remains a challenge. Perhaps the best example of this is given by our analysis of a fourth Sima de los Huesos sample, Femur XIII, for which a minimum sequence length cutoff of 46 bp must be applied to ensure that the fraction of spurious alignments is restricted to less than 1%. This result shows that microbial contamination is so abundant in this sample that the commonly used cutoffs of 35 bp length or shorter (Table S1) is insufficient to reduce the effect of spurious alignments to conservative levels. As more data from highly degraded material become available, it will be crucial to ensure that spurious alignments are quantified to avoid false results.

On a broader level, our results show that the genetic analysis of poorly preserved ancient biological material is not only limited by our ability to extract and sequence the DNA it may contain, but also by our ability to distinguish sequences that are endogenous to the organism from the overwhelming majority of microbial contamination. Molecular methods have been developed in the past to decrease the fraction of microbial contamination. These methods used restriction enzymes that cut motifs occurring preferentially in contaminant DNA [16], enriched for endogenous DNA fragments via hybridization capture [28] or depleted contaminant DNA prior to DNA extraction [29, 30]. Further research will be needed to establish how these methods can contribute to the study of highly-degraded samples.

We conclude that while spurious alignments are an inevitable issue for the analysis of short ancient sequences, their influence can be accurately assessed and limited by appropriate filtering. Together with further refinement of molecular methods our approach paves the way towards the study of older or more degraded samples.

## Materials and methods

### Modifying the human reference genome

The human reference genome (hg19/GRCh37) was used as a template to create a genome with additional single nucleotide changes. These changes were introduced in conserved regions where the reference human base is identical to the aligned bases of the chimpanzee pantro4 genome, the high-coverage genomes of Altai Neandertal [21] and Denisova [20], 24 high-coverage modern human genomes [20, 31], and all 2,504 modern human individuals of the 1000 Genomes Project data phase 3 [32]. Sites five base-pairs up- and down-stream of all indels detected in these datasets were excluded. Sites were also required to fall outside of simple repeats annotated using the Tandem Repeat Finder [33] and to overlap positions of unique mapability based on 35mers [21]. Bases were changed every 100 bp. If a change fell in a region that was excluded, the closest included position was determined and chosen as new location if it was at least 75 base-pairs from the closest adjacent changed site. Bases were replaced by other bases so that the overall nucleotide composition was kept identical to that of the hg19 genome. A total of 18,002,060 sites were modified.

### Sequence Data and Alignments to the modified reference

We used one lane of Illumina HiSeq 2500 sequencing data from the Mezmaiskaya1 Neandertal individual (library R5661; see Suppl. 2 in ref. [22]) and the published sequences from Sima de los Huesos samples [34] Femur fragment, Incisor, Molar and FemurXIII. Both datasets were generated with the same extraction method [3] and the single stranded DNA library protocol [35].

As negative control – i.e. as a sample for which we do not expect to see any spurious alignments – we used modern human DNA that was sheared to short fragments of similar size to those in ancient samples. In details, DNA was extracted from the blood of a healthy human donor using the Gentra Puregene Blood Kit (Qiagen). One microgram of DNA was sheared for 2h using the Covaris S2 ultrasonicator (shearing parameters: intensity 5, cycler/burst 1000, duty cycles 10%) to obtain a fragment size distribution that mimics that of ancient DNA. A 200 ng aliquot of sheared DNA was then used as input for silica-based DNA extraction [3]. A single-stranded library [35] was prepared from 2.5 μl of the resulting DNA extract (5% of the extract). The library was amplified using Accuprime Pfx DNA polymerase (Thermo Fisher Scientific) [36] and a pair of indexing primers containing a sample-specific combination of 7 bp indices [37]. The indexed library was sequenced on 6 lanes of a HiSeq 2000 (Illumina) in 2x 76 bp paired-end configuration with two index reads [37]. Sequences without perfect matches to the expected index combination were discarded.

For our positive control – i.e. a sample with solely spurious alignments – we used 3,860 bacteria genomes from the European Nucleotide Archive listed here http://www.ebi.ac.uk/genomes/bacteria.details.txt. The genomes were then fragmented to 32 bp, an arbitrary length shorter than 35 bp, using a one bp tiling, resulting in a total of ~7.2 billion unique sequences. Ambiguous bases were replaced with one randomly chosen representative base. All sequence data were mapped to the modified human reference genome using *bwa* [38] with options ‘-n 0.01 –o 2 –l 16500’ matching those used for the ancient samples [14, 20]. Sequences were merged when they appeared to originate from a PCR duplicate by means of *bam-rmdup* (https://bitbucket.org/ustenzel/biohazard-tools). Paired-end sequences and sequences shorter than 20 bp were disregarded.

### Length-dependent mappability tracks

We used the software *gem* [39] to generate maps of unique mappability of different lengths for the human reference genome (GCRh37/hg19) including decoy sequences [32]. The program was run for lengths of 20, 23, 26, 29, 32 and 35 base-pairs allowing for up to one mismatch in alignments. To determine whether a sequence was mappable, we first chose the largest mappability track that was not longer than the sequence length. The sequence was deemed uniquely aligned if it contained a uniquely mappable motif in the reference within its alignment. All analyses involve this filtering.

### Features of spurious alignments

Sequences that mapped to the modified reference genome and overlap mutated sites were used to determine characteristics of spurious and true alignments. Alignments were classified as true if they showed the human reference base and as spurious if they showed the mutated variant or any other allele than the human reference. For both spurious and true alignments, we calculated:

1. The proportion of mismatches, i.e. the number of observed mismatches relative to the modified reference genome divided by sequence length. For sequences that did not match the modified reference’s allele, we subtracted one mismatch to compensate for the mismatch caused by the artificially mutated site. This correction was applied to true and spurious alignments, alike.
2. The number of insertion and deletions (indels), extracted from the CIGAR field in the bam/sam format files.
3. The patterns of nucleotide substitutions, determined by comparing the sequences to the unmodified hg19 reference.

To minimize the impact of cytosine deamination we make use of the preserved strand orientation of sequences prepared with the single-stranded library protocol [35] and disregarded alignments in the forward orientation if either the mutated or original human reference state was C, or alignments in reverse orientation if the mutated or original human reference state was G. Due to this filter 37% of mutated sites (C-to-G or G-to-C) are disregarded.

### Quantifying the fraction of spurious alignments

To calculate the proportion of spurious alignments we make use of the number of alignments classified as truly related (*N_T_*) and the number of alignments classified as spurious (*N_¬T_*) as described in the previous section. A small fraction of spurious alignments is expected to show the human reference base by chance. To correct for this, we assume that all spurious alignments contain the maximum number of mismatches. A spurious alignment is then misclassified with the probability of 1/3 of the mismatch proportion along the sequence, and we conservatively correct the *N_T_* and *N_¬T_* counts to compensate for spurious misclassified alignments by calculating

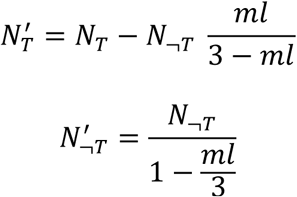

where *ml* denotes the proportion of mismatches allowed at a given length in the mapping procedure. With these corrected counts, we calculate the spurious alignment proportion as:

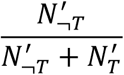

### Lineage assignment

Informative sites were determined by sampling one random allele from each of the genotypes of a modern human (Mbuti), the Altai Neandertal and the Denisovan genomes after applying the minimum set of filters described in [21]. To call the ancestral state at each site we used whole genome alignments of five primates (chimpanzee, bonobo, gorilla, orangutan, and macaque), and required that at least four of them agree. Derived sites were assigned to the following four lineages: Modern Human, Neandertal, Denisovan, Neandertal-Denisovan.

For each dataset, we iterated over all sequences and calculated the percentage of derived alleles of each class that are shared. All T within the last three terminal positions of sequences were disregarded to minimize the impact of C-to-T changes due to deamination.

## Funding

This study was supported by the Max Planck Society, funded by Max Planck Foundation grant “No. 31-12LMP Pääbo” and by the European Research Council grant No. 694707 to Svante Pääbo.

## Authors’ contributions

MM and KP designed the study. All authors analyzed the data, interpreted the results, wrote the manuscript the final manuscript.

## Acknowledgments

We are thankful to Michael Dannemann, Janet Kelso, Fabrizio Mafessoni, Svante Pääbo, Udo Stenzel, and the Neandertal and Bioinformatics groups of the Evolutionary Genetics department for helpful discussions and suggestions during the development of the project. We are also grateful to Marie Gansauge and Birgit Nickel for the extraction and library preparation of the modern human sample.

